# Plasticity of cell migration resulting from mechanochemical coupling

**DOI:** 10.1101/644880

**Authors:** Yuansheng Cao, Elisabeth Ghabache, Wouter-Jan Rappel

## Abstract

Eukaryotic cells can migrate using different modes, ranging from amoeboid-like, during which actin filled protrusions come and go, to keratocyte-like, characterized by a stable morphology and persistent motion. How cells can switch between these modes is still not well understood but waves of signaling events on the cell cortex are thought to play an important role in these transitions. Here we present a simple two component biochemical reaction-diffusion model based on relaxation oscillators and couple this to a model for the mechanics of cell deformations. Different migration modes, including amoeboid-like and keratocyte-like, naturally emerge through phase transitions determined by interactions between biochemical traveling waves, cell mechanics and morphology. The model predictions are explicitly verified by systematically reducing the protrusive force of the actin network in experiments using wild-type *Dictyostelium discoideum* cells. Our results indicate the importance of coupling signaling events to cell mechanics and morphology and may be applicable in a wide variety of cell motility systems.

## Introduction

Eukaryotic cell migration is a fundamental biological process that is essential in development and wound healing and plays a critical role in pathological diseases, including inflammation and cancer metastasis Ridley et al. (2003); Roussos et al. (2011); Montell (2003). Cells can migrate using a variety of modes with a range of corresponding morphologies. The repeated extensions and retractions of pseudopods in amoeboid-like cells, for example, result in a constantly changing morphology and random migration while keratocyte-like cells have a stable and broad actin-rich front, a near-constant shape, and move in a persistent fashion Webb and Horwitz (2003); Keren et al. (2008). Furthermore, many cells do not have a unique migration mode and can switch between them, either as a function of the extracellular environment or upon the introduction of a stimulus Paul et al. (2017); Bergert et al. (2012); Charras and Sahai (2014); Liu et al. (2015); Petrie and Yamada (2016); Miao et al. (2017). This plasticity is currently poorly understood and is thought to play a role in pathological and physiological processes that involve cell migration, including cancer metastasis Friedl and Alexander (2011).

A key step in cell migration is the establishment of an asymmetric and polarized intra-cellular organization where distinct subsets of signaling molecules, including PAR proteins, Rho family GTPases and phosphoinositides, become localized at the front or back of the cell Jilkine and Edelstein-Keshet (2011); Rameh and Cantley (1999); Goldstein and Macara (2007); Raftopoulou and Hall (2004); Rappel and Edelstein-Keshet (2017). In the absence of directional cues, this symmetry breaking can be a spontaneous and dynamic process with waves of cytoskeletal and signaling components present on the cell cortex Vicker (2002); Weiner et al. (2007); Whitelam et al. (2009); Case and Waterman (2011); Gerisch et al. (2012); Allard and Mogilner (2013); Gerhardt et al. (2014); Barnhart et al. (2017). Addressing this spontaneous symmetry breaking and the role of waves have generated numerous theoretical studies Jilkine and Edelstein-Keshet (2011); Meinhardt (1999); Otsuji et al. (2007); Csikász-Nagy et al. (2008); Beta et al. (2008); Mori et al. (2008); Xiong et al. (2010); Knoch et al. (2014); Miao et al. (2017). Most models, however, study cell polarity in the context of biochemical signaling and do not consider cell movement or deformations originated from cell mechanics. This may be relevant for nonmotile cells including yeast Park and Bi (2007); Slaughter et al. (2009) but might not be appropriate for motile cells where the coupling between intracellular pathways and cell shape can be crucial in determining the mode of migration Camley et al. (2013, 2017). Furthermore, most of these models only focus on one specific migration mode and do not address transitions between them. Therefore, it remains an open question how cell mechanics, coupled to a biochemical signaling module, can affect spontaneous cell polarity and can determine transitions between cell migration modes.

Here we propose a novel model that couples an oscillatory biochemical module to cell mechanics. Our choice of the biochemical model was motivated by recent findings that the self-organized phosphatidylinositol (PtdIns) phosphate waves on the membrane of Dictyostelium cells exhibit characteristics of a relaxation oscillator Arai et al. (2010). Our model is able to generate amoeboid-like, keratocyte-like, and oscillatory motion by varying a single mechanical parameter, the protrusive strength, without altering the biochemical signaling pathway. We determine how the transitions depend on these parameters and we show that keratocyte-like motion is driven by an emergent traveling wave whose stability is determined by the mechanical properties of the cell. Finally, we experimentally obtain all three migration modes in wild-type *Dictyostelium discoideum* and explicitly verify model predictions by reducing the actin protrusive force using the drug latrunculin B. Our model provides a unified framework to understand the relationship between cell polarity, motility and morphology determined by cellular signaling and mechanics.

## Models and results

### Model

Our two-dimensional model is composed of two modules: a biochemical module describing the dynamics of an activator-inhibitor system which works in the relaxation oscillation regime, and a mechanical module that describes the forces responsible for cell motion and shape changes (Fig.1a). Our biochemical module consists of a reaction-diffusion system with an activator *A* (which can be thought of as PtdIns phosphates and thus newly-polymerized actin Gerhardt et al. (2014); Miao et al. (2019)) and an inhibitor *R* (which can be thought of as the phosphatase PTEN). This activator and inhibitor diffuse in the cell and obey equations that reproduce the characteristic relaxation oscillation dynamics in the PtdIns lipid system Arai et al. (2010):

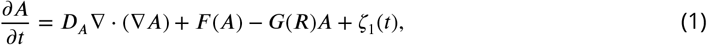

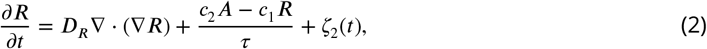

where *D_A_* and *D_R_* are the diffusion coefficients for *A* and *R*, respectively. In these expressions, *F*(*A*) is the self-activation of the activator with a functional form that is similar to previous studies: 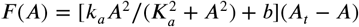 Miao et al. (2017). The activator is inhibited by *R* through the negative feedback *G*(*R*) = *d*_1_ + *d*_2_*R* while *R* is linearly activated by *A* (see Methods and Materials). The timescale of the inhibitor *τ* is taken to be much larger than the timescale of the activator. Finally, to ensure robustness to stochasticity, we add uniformly distributed spatial white noise terms 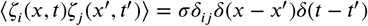.

**Figure 1.**
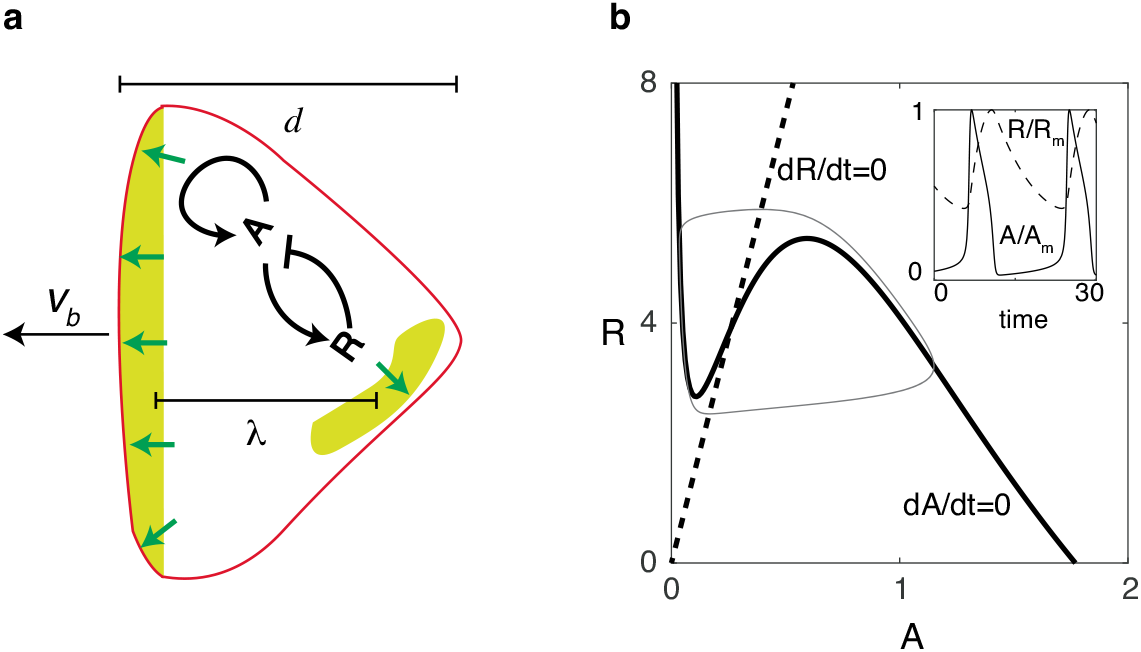
Reaction diffusion model coupled to a mechanical model. (a) Schematic illustration of the two-dimensional model: a self-activating activator field *A*, indicated in color, drives the movement of the cell membrane through protrusive forces that are normal to the membrane (green arrows). One successive wave is generated behind the original one after a distance of *λ*. The cell’s front-back distance is *d*, and the cell boundary is pushed outward with speed *υ_b_*. (b) Nullclines of the activator (solid line) and inhibitor (dashed line), along with the resulting trajectory in phase space (gray thin line). The inset shows the oscillations of *A* and *R*, normalized by their maximum values *A_m_* and *R_m_*.

Nullclines for this system are shown in Fig.1b, where we have chosen parameters such that the fixed point is unstable and the system operates in the oscillatory regime. As a result of the separation of timescales for *A* and *R*, the dynamics of *A* and *R* are characteristic of a relaxation oscillator (inset of Fig.1b): *A* reaches its maximum quickly, followed by a much slower relaxation phase.

To generate cell motion, we couple the output of the biochemical model to a mechanical module which incorporates membrane tension and bending as well as protrusive forces that are proportional to the levels of activator *A* and normal to the membrane, similar to previous studies Shao et al. (2010, 2012) (see Methods and Fig.1a). To accurately capture the deformation of the cell in simulations, we use the phase field method Shao et al. (2010); Ziebert et al. (2011); Shao et al. (2012); Najem and Grant (2013); Marth and Voigt (2014); Camley et al. (2017); Cao et al. (2019). Here, an auxiliary field *ϕ* is introduced to distinguish between the cell interior (*ϕ* = 1) and exterior (*ϕ* = 0), and the membrane can be efficiently tracked by the contour *ϕ* =1/2. Coupling this field to the reaction-diffusion equations can guarantee that no-flux boundary conditions at the membrane are automatically satisfied Kockelkoren et al. (2003). The evolution of the phase-field is then determined by the force balance equation:

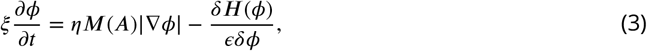

where *ξ* is a friction coefficient, *ϵ* is the boundary width of the phase field, and *H*(*ϕ*) is a Hamiltonian energy including the membrane tension, parameterized by *γ*, bending energy, and area conservation (see Methods and Supplementary Information). The first term on the right hand side describes the actin protrusive force, parameterized by *η*, and acts on the cell boundary since |∇*ϕ*| is non-zero only in a region with width *ϵ*. *M* formulates the dependence of the protrusive force on the activator levels and is taken to be sigmoidal: 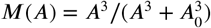. As initial conditions, we use a disk with radius *r* with area *S* = *πr*^2^ and set *A* = *R* = 0. Default parameter values for our model are estimated from experimental data and given in Table 1.

**Table 1.** Model Parameters

| Parameter | Description | Value |
| --- | --- | --- |
| $\gamma$ | Tension | 2 pN |
| $\kappa$ | Bending | 2 pN $\mu\text{m}^2$ |
| $\epsilon$ | Width of phase field | 2 $\mu\text{m}$ |
| $B_S$ | Cell area conservation strength | 10 pN/$\mu\text{m}$ |
| $\xi$ | Friction coefficient | 10 Pa s/$\mu\text{m}$ |
| $k_a$ | Activation rate | 10 $\text{s}^{-1}$ |
| $K_a$ | Activation threshold | 1 $\mu\text{M}$ |
| $b$ | Basal activation rate | 0.1 $\mu\text{M} \text{s}^{-1}$ |
| $A_t$ | Total activator concentration | 2 $\mu\text{M}$ |
| $d_1$ | Basal degradation rate | 1 $\text{s}^{-1}$ |
| $d_2$ | Degradation rate from inhibitor | 1 $\mu\text{M}^{-1} \text{s}^{-1}$ |
| $c_1$ | Inhibitor degradation coefficient | 1 |
| $c_2$ | Inhibitor activation coefficient | 15 |
| $\tau$ | Time scale of negative feedback | 10 s |
| $D_A$ | Activator diffusion coefficient | 0.5 $\mu\text{m}^2/\text{s}$ |
| $D_R$ | Inhibitor diffusion coefficient | 0.5 $\mu\text{m}^2/\text{s}$ |
| $\sigma$ | Noise intensity | 0.01 $\mu\text{M}^2/\text{s}$ |
| $\Delta t$ | Time step | 0.001 s |
| $n, m$ | Space grid size | 256,256 |
| $L_{x,y}$ | Space size | 50,50 $\mu\text{m}$ |

### Computational results

We first examine the possible migration modes as a function of the protrusive strength *η* for fixed area *S*, parametrized by the radius *r* of the disk used as initial condition, and default parameters. As shown in Fig.2, there are three distinct cell migration modes. When *η* is small, activator waves initiate in the interior and propagate to the cell boundary. However, the protrusive force is too small to cause significant membrane displacement. Consequently, the cell is almost non-motile and the activator and inhibitor field show oscillatory behavior (Fig.2a I and Movie S1).

**Figure 2.**
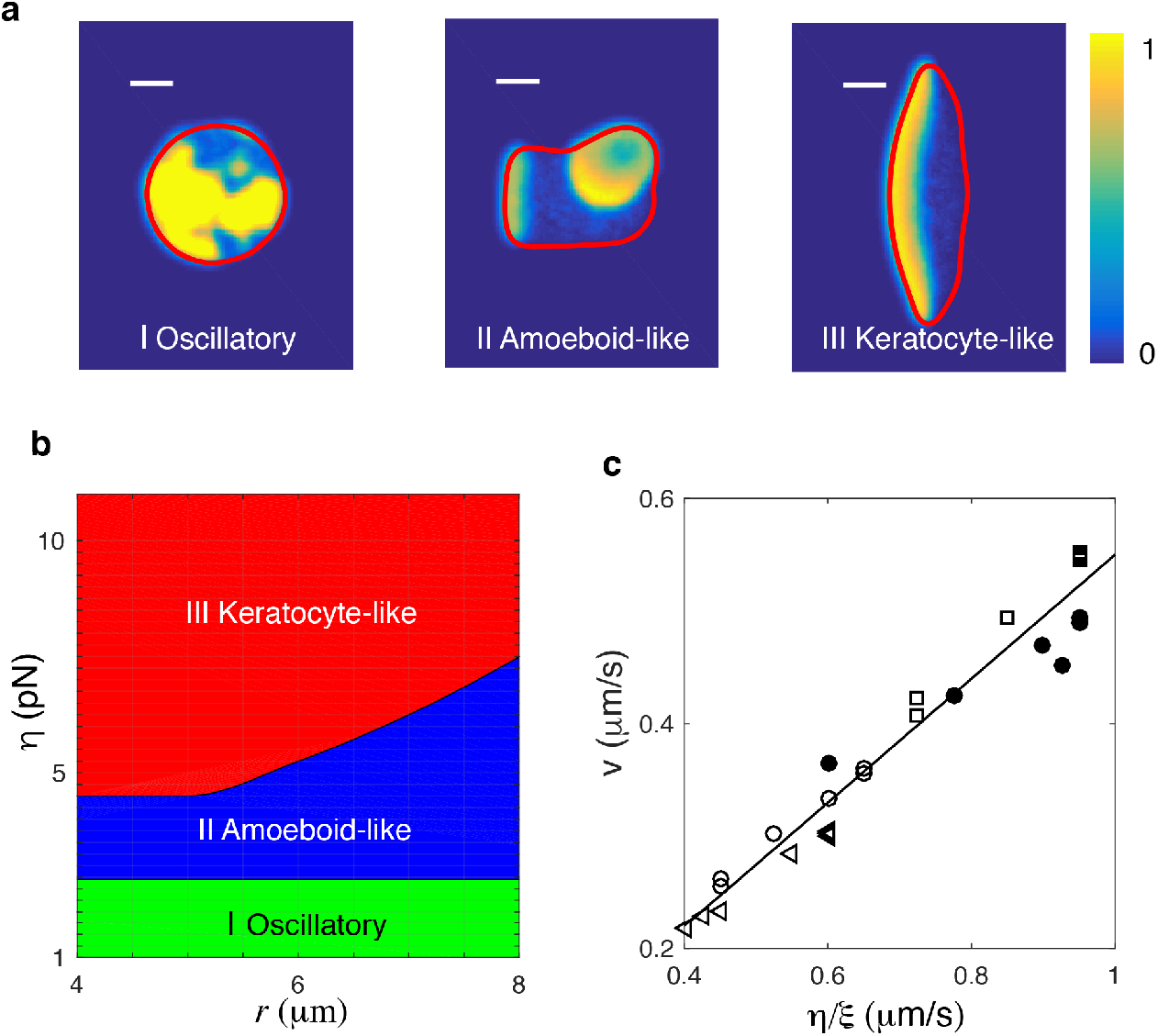
Different cell migration modes can be captured in the model by varying the protrusive strength *η*. Snapshots of a simulated cell showing (I) an oscillatory cell (*η* = 2 pN), (II) an amoeboid-like cell (*η* = 4 pN), and (III) a keratocyte-like cell (*η* = 11 pN). All other parameters were assigned the default values and *r* = 8*μ*m. Here, the activator concentration is shown using the color scale and the cell membrane is plotted as a red line (scale bar 5*μ*m). (b). Phase diagram determined by systematically varying *η* and the initial radius of the cell, *r*. (c). The speed of the keratocyte-like cell as a function of *η*/*ξ*. The black line is the predicted cell speed with *υ_b_* = *αη*/*ξ*, where *α* ≈ 0.55. Symbols represent simulations using different parameter variations: empty circles, default parameters;triangles, *ξ* = 2*ξ*_0_; filled circles, *γ* = 2*γ*_0_; squares, *τ* = *τ*_0_/2. **Figure 2–Figure supplement 1.** Speed of keratocyte-like cells as a function of the surface tension and timescale of the inhibitor **Figure 2–Figure supplement 2.** Transition from oscillatory cells to amoeboid-like cells **Figure 2–Figure supplement 3.** Effects of tension **Figure 2–Figure supplement 4.** Parameter variations in the model **Figure 2–Figure supplement 5.** Oscillatory cells for strong and weak area conservation **Figure 2–Figure supplement 6.** Excitable dynamics can reproduce identical qualitative results

As *η* increases, an activator wave that reaches the boundary can create membrane deformations, leading to the breaking of spatial homogeneity. This wave, however, is competing with other traveling waves that emerge from random positions. Consequently, the cell exhibits transient polarity, moves in constantly changing directions, and displays amoeboid-like migration (Fig.2a II and Movie S2). When *η* is increased further, protrusions generated by activator waves reaching the cell boundary become even larger. As a result of the coupling between the waves and membrane mechanics, a single traveling wave will emerge within the cell, characterized by a broad and stationary band of high levels of activator. This wave pushes the membrane forward in a persistent direction with constant speed and the cell will adopt a steady keratocyte-like morphology, even in the presence of noise (Fig.2a III and Movie S3).

The transition from oscillatory dynamics to amoeboid-like unstable cell motion can be understood by considering the coupling between the traveling waves and membrane motion. In the Supplementary Information we show that these traveling waves, that emerge naturally in systems of relaxation oscillators Kopell and Howard (1973); Keener (1980); Murray (2002) are stable as long as the activator front can “outrun” the inhibitor’s spreading speed. This condition results in a minimal wave speed *c*_min_ that depends on *D_R_* and *τ* (see Methods and Materials). The activator wave pushes the membrane outward and will keep propagating as long as the boundary can keep up with the wave speed. In our model, the membrane is pushed outward by a protrusive force resulting in a speed approximately given by *υ_b_* ~ *αη*/*ξ*, where *α* is the boundary-averaged value of *M*(*A*), that depends on mechanical parameters and is independent of biochemical parameters (see Methods and Materials and Fig2–Figure Supplement 1). This implies that if the speed of the membrane is less that the minimum speed of the activator wave (*υ_b_* < *c*_min_) there will be no significant membrane motion. On the other hand, when *υ_b_* > *c*_min_, a traveling wave can be selected by matching the wave speed and the boundary speed:

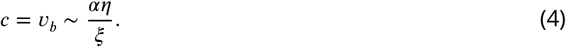

The above equation indicates that when *η* > *ξc*_min_/*α*, the cell will break its symmetric shape through traveling waves that deform the membrane. In simulations, the critical value of *η*, *η*_*c*,1_, for which the oscillating cell becomes amoeboid-like can be determined by slowly increasing *η* and computing the center-of-mass speed of the cell. These simulations show that the bifurcation between the two migration modes is subcritical (Fig2–Figure Supplement 2). Furthermore, these simulations reveal that, as argued above, *η*_*c*,1_ depends on both *D_R_* and *τ* (Fig2–Figure Supplement 2).

Once a traveling wave is able to generate membrane deformations, why does it not always result in stable, keratocyte-like motion with a single traveling wave in the cell’s interior? Notice that in our model, if the spatial extent between the cell front and the back, *d*, is larger than the wavelength of the activator wave *λ* (schematically shown in Fig.1 a), a new wave will be generated behind the original one. This wavelength can be approximated by 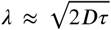 (where we have taken *D_A_* = *D_R_* = *D* for simplicity) such that stable keratocyte-like cells driven by a single wave are only possible when *d* < *λ*. This front-back distance, however, depends on the balance between protrusive force and membrane tension at the traveling wave’s lateral ends. Increasing values of *η* result in a broader front and therefore smaller values of *d* (see Methods and Materials for a discussion of tension effects and Fig2–Figure Supplement 3). As a consequence, as *η* is increased, waves propagating within the cell eventually become stable, resulting in a single, propagating wave and a cell with a keratocyte-like morphology. This also implies that, for equal values of the protrusive strength, the polarity of larger cells will be less stable than the polarity of smaller ones.

From the above analysis, it becomes clear that the protrusive strength and size of initial disk, and thus cell area, are critical parameters in determining the stability of the polarity established by interactions between traveling waves and moving boundaries. In simulations, we therefore determine the phase diagram in the (*η*, *r*) space by systematically varying the size (with step size 0.5 *μ*m) and protrusive force (with step size 0.25 pN) while keeping all other parameters fixed (Fig.2b). We constrain our cell area to be within the physiological relevant ranges with a maximum initial cell size of *r* = 8*μ*m, corresponding to an area *S* ~ 200*μ*m^2^ (see Methods and Materials and Fig2–Figure Supplement 4 for an extension of the phase space to larger values of *r*). As stated above, there are three distinct phases, corresponding to the three different cell migration modes of Fig.2a. The transition from oscillatory to amoeboid-like motility occurs at small *η* and is independent of cell size, as predicted in Eq.(4). The transition from unstable to stable polarity, and therefore keratocyte-like motion, occurs for a critical value of *η*, *η*_*c*,2_, which increases for increasing values of *r*. The latter transition depends on parameters that affect either *d* or *λ* (Fig2–Figure Supplement 4). Finally, we find the speed of the keratocyte-like cell is linearly dependent on *η*/*ξ*, and independent of other parameters, as predicted by Eq. 4 (Fig.2c).

In summary, our model predicts that a sufficient decrease of *η* can destabilize keratocyte-like cells, resulting in cells that employ amoeboid-like migration, and can transform keratocyte-like and unstable cells into oscillatory cells. Furthermore, for decreasing protrusive force, the keratocyte-like cells should have a reduced speed and cell size.

### Experimental Results

To test our model predictions, we carry out experiments using wild-type Dictyostelium cells (see Methods and Materials). In vegetative food-rich conditions, during which food is plentiful, most cells migrate randomly using amoeboid-like motion. Starvation triggers cell-cell signaling after which cells become elongated and perform chemotaxis. However, we found that starving cells for 6h at sufficiently low density is enough to prevent cell-cell signaling. Under these conditions, the majority of cells still moves as amoeboid-like cells but a significant fraction, approximately 20-50%, migrates in a keratocyte-like fashion (Movie S4). These cells adopt a fan-shaped morphology and move unidirectionally, as was also observed in certain Dictyostelium mutants Asano et al. (2004). Employing these low density conditions, we can alter the cell’s protrusive force by interrupting actin polymerization using the drug latrunculin B, an inhibitor of actin activity. Our model predicts that as the concentration of latrunculin increases, keratocyte-like cells are more likely to switch to unstable or oscillatory cells. Furthermore, the size and speed of the remaining keratocyte-like cells should decrease.

A snapshot of starved cells is shown in Fig. 3a, before (left panel) and after exposure to latrunculin (right panel). Higher magnification plots of the amoeboid-like cells (top two panels) and of a keratocyte-like cell are shown to the right. In these panels, the actin distribution is visualized with the fluorescent marker limE-GFP. As can be seen by comparing the snapshots in Fig. 3a, the number of keratocyte-like cells decreases after the exposure of latrunculin. This decrease is due to keratocyte-like cells becoming unstable and switching to the amoeboid-like mode of migration. We quantify the percentage of keratocyte-like cells, as well as the speed and shape, as a function of time for different concentrations of latrunculin for at least 100 cells. As shown in Fig. 3b, the percentage of keratocyte-like cells decreases upon the introduction of latrunculin. Furthermore, this decrease becomes more pronounced as the concentration of latrunculin is increased (inset Fig. 3b), consistent with our model prediction.

**Figure 3.**
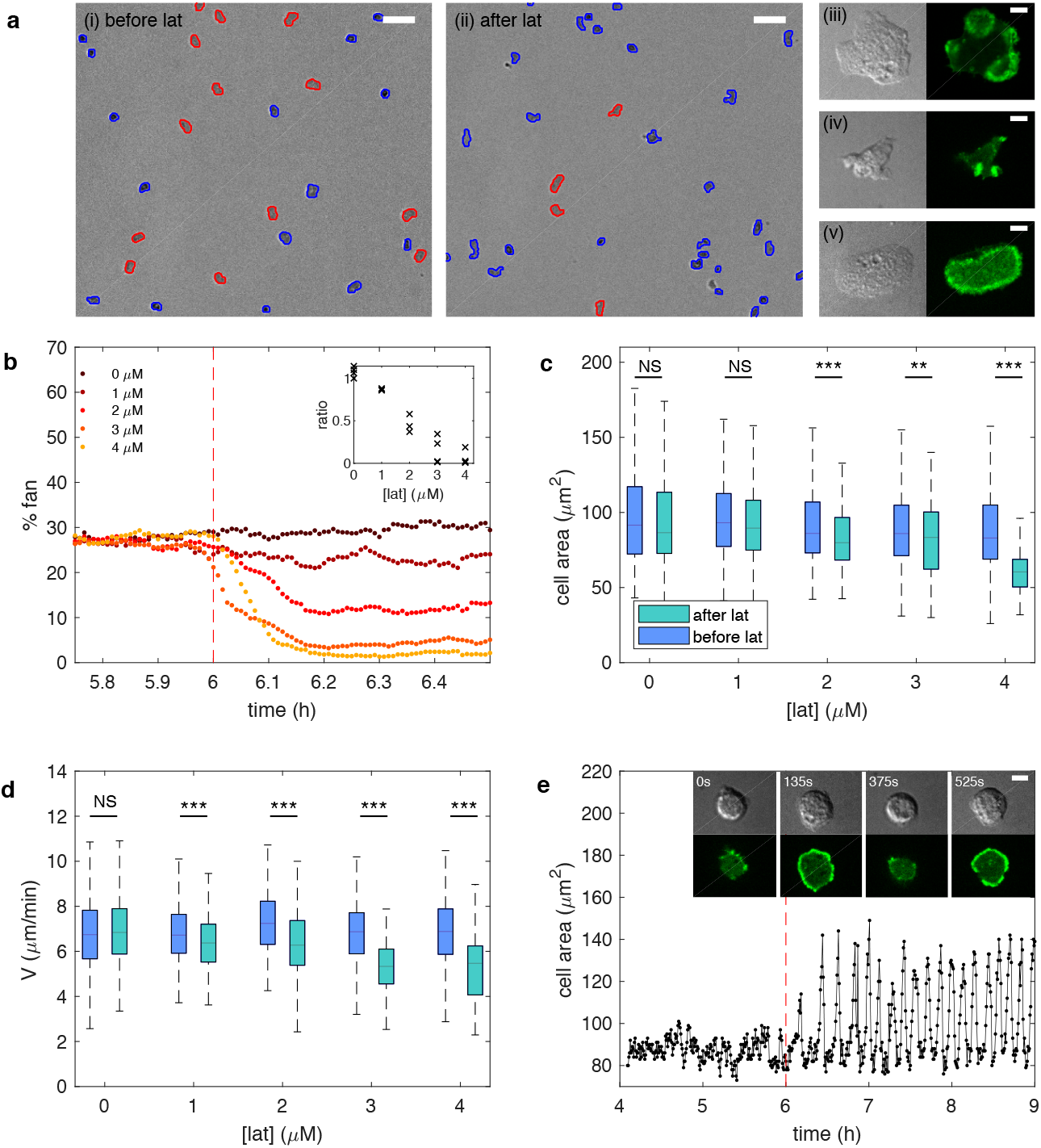
Experiments reveal different migration modes in Dictyostelium cells. (a) Snapshot of starved Dictyostelium cells before (left) and after (right) exposure to latrunculin B. Amoeboid-like cells are outlined in blue while keratocyte-like cells are outlined in red (scale bar: 50 *μ*m). The panels on the right show high magnification views of amoeboid-like (top two) and keratocyte-like cells in which the freshly polymerized actin is visualized with limE-GFP (scale bar: 5 *μ*m). (b) Percentage of keratocyte-like cells as a function of time for different concentrations of latrunculin B (introduced at 6 h, dashed line). Inset shows the ratio of keratocyte-like to all cells as a function of the latrunculin concentration for three repeats. (c) The cell area size of keratocyte-like cells before and after latrunculin exposure as a function of concentration. (d) The speed of keratocyte-like cells before and after latrunculin exposure as a function of concentration. (e) Basal cell area as a function of time for a cell that transitioned from amoeboid-like to oscillatory. Insets show snapshots of the cell at different time points (scale bar: 5 *μ*m). **Figure 3–Figure supplement 1.** The front-back distance of keratocyte-like cells before and after latrunculin exposure **Figure 3–Figure supplement 2.** The average period and coefficient of variation for area oscillations in oscillating cells after the exposure to latrunculin

To further verify the model predictions, we quantify the cell area size *S* and cell speed for the keratocyte-like cells. Both the size (fig.3c), front-back distance (Fig3–Figure Supplement 1) and speed (fig.3d) decrease after the introduction of latrunculin. This decrease becomes more significant for larger concentrations of latrunculin, again consistent with our predictions.

Finally, our model predicts that, with a relaxed area conservation, a sufficient reduction of protrusive strength results in the the appearance of oscillatory cells with oscillating basal area size (Fig2–Figure Supplement 5). Indeed, after the exposure to latrunculin, a small fraction of cells are observed to display oscillatory behavior characterized by repeated cycle of spreading and contraction, resulting in a basal surface area that oscillates, similar to the engineered oscillatory cells of Miao et al. (2017) (Fig. 3e). Cell tracking reveals that these cells originate through a transition from the unstable, amoeboid-like state to the oscillatory state. Interestingly, the observed oscillation in surface area is often very regular (Fig. 3e). We find that the average period of different cells is largely independent of the latrunculin concentration, ranging from 6.2 ± 0.9 min for 1 *μ*M to 6.6 ± 0.9 min for 4 *μ*M, while the coefficient of variation, defined as the ratio of the standard deviation and the mean, for single cell periods varies between 0.31 ± 0.12 min (1 *μ*M) and 0.20 ± 0.10 min (4 *μ*M; Fig3–Figure Supplement 2). In conclusion, our experimental results are in full agreement with the model predictions.

## Discussion

In this paper, we propose a simple but unified paradigm to understand cell migration and cell morphology. As in previous modeling studies Miao et al. (2017, 2019), our model displays three different migration modes which can be induced by varying the protrusive force. This is attractive since the switching of these modes can occur on a timescale that is shorter than gene expression timescales (Fig. 3b), suggesting that a single model with conserved components should be able to capture all three modes. Importantly, our model predictions are verified in experiments using wild-type Dictyostelium cells which, under our conditions, exhibit all three migration modes. These experiments show that upon the introduction of latrunculin the speed of keratocyte-like cells decrease. This is perhaps not surprising since latrunculin inhibits actin polymerization which can be expected to result in smaller cell speeds. Our experiments also show, however, that the area of the moving keratocyte-like cells decreases in the presence of latrunculin. In addition, and more importantly, our experiments demonstrate that transitions between the migration modes can be brought about by reducing the protrusive strength of the actin network. These non-trivial effects of latrunculin are consistent with the predictions of our model.

Key in our model is the coupling of traveling waves generated through biochemical signaling and cell mechanics. Our main finding is that cell migration is driven by the traveling waves and that persistent propagation of these waves result in keratocyte-like cells with a broad and stable front. This stable front is only present if the front-back distance is smaller than the biochemical wavelength. Reducing the protrusive strength results in a larger front-back distance, resulting in unstable, amoeboid-like migration. For even smaller values of the protrusive strength, cells display oscillatory behavior. For these values, the membrane speed is smaller than the minimum biochemical wave speed.

Recent studies by Miao et al. (2017) have proposed a model that can generate all three migration modes observed in experiments of engineered Dictyostelium cells Miao et al. (2017, 2019). Our current model is distinct from these studies in several ways. First, the migration mode transitions in our model are induced by the mechanical module with the same biochemical components, while in Miao et al. (2017) the transitions are generated by changing the dynamics of the biochemical signaling pathways from, e.g., excitable to oscillatory. In addition, the biochemical module in our model is much simpler and only contains an activator and inhibitor while the model of Miao et al. (2017) requires additional feedback from a postulated polarity module. Of course, our work does not exclude the existence of this polarity module but it shows that we can explain the observed cell morphologies and movement within a minimal framework of coupling two biochemical components and cell mechanics. Second, based on the earlier measurements of Arai et al. (2010), our model assumes that the biochemical module operates as a relaxation oscillator instead of nested excitable networks Hecht et al. (2011); Miao et al. (2017, 2019). Note however, that in our model we can tune the negative feedback to make the biochemical module operate in the excitable regime. We have explicitly verified that qualitatively similar migration modes and transitions are observed if our model is excitable (Fig2–Figure Supplement 6). In the excitable version of our model, however, the oscillations in the non-motile mode are, in general, less regular and periodic than the ones obtained in the relaxation oscillator version. From the statistical features of the periods obtained from oscillatory cells in experiments, it is likely that the cellular signaling dynamics can be most accurately described by relaxation oscillation models. As a final distinction, we point out that the biochemical and mechanical module in the model of Miao et al. (2017) are solved separately on a 1D ring. As a result, the keratocyte-like cells display constant excitations at the front that travel along the membrane in the lateral direction rather than stationary activator bands, as observed in the experiments and in our model.

Several future extensions of our study are possible. First, our current study is restricted to two dimensional geometries while actual cell motion is of course three dimensional. Extending our model to 3D would allow us to relax the area conservation constraint and should result in cells for which the basal surface area show clear oscillations that are coupled to extensions away from the surface (Fig2–Figure Supplement 5). Second, it should be possible to couple the biochemical model to an upstream chemotaxis pathway, allowing it to address directed motion. Third, it would be interesting to compare wave dynamics obtained in our model with waves observed in giant Dictyostelium cells Gerhardt et al. (2014). In addition, alternative biochemical models in which parameters determine the qualitatively different dynamics can be studied Miao et al. (2017). Furthermore, our study predictions may also be verified in other cell types. For example, we predict that overexpression of actin in fast moving cells should result in cells migrating with keratocyte-like morphologies while disturbing actin polymerization in keratocytes could lead to unstable migration. Finally, it would be interesting to determine how the feedback between mechanical and biochemical modules can potentially help understand other cell migration processes.

## Acknowledgments

We thank Brian A. Camley for many useful discussion. This work was supported by the National Science Foundation under grant PHY-1707637 and HFSP number LT000371/2017-C.

## Methods and Materials

### Full model

Our model for the cell boundary and cell motion is detailed in earlier studies Shao et al. (2010, 2012); Camley et al. (2014). Briefly, we model the cell boundary as an interface with tension and bending, driven in this study by activator *A* at the front. Cell motion obeys the overdamped force balance equation ***F**_act_* + ***F**_mem_* + ***F**_area_* + ***F**_fric_* = 0 where ***F**_act_* is the active force proportional to the activator concentration; ***F**_mem_* describes the membrane tension and bending forces; ***F**_area_* represents area conservation to prevent cells from expanding or shrinking indefinitely and ***F**_fric_* is a friction force. The active force from the activator is governed by 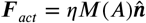, where 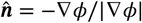 is the outward-pointing normal direction of the membrane, and 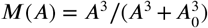 where *A*_0_ represents a threshold value for activation of protrusive force. The membrane tension and bending forces are computed using the functional derivative Camley et al. (2014) 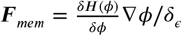, with 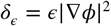 and

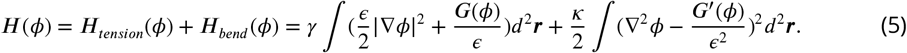

Here, *G*(*ϕ*) is a double well potential with minima at *ϕ* = 1 and *ϕ* = 0: *G*(*ϕ*) = 18*ϕ*^2^(1 - *ϕ*)^2^. We implement area conservation as 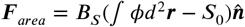 where *B_S_* represents the strength of the area conservation and *S*_0_ is the prescribed area size determined by initial cell radius *r* through *S*_0_ = *πr*^2^. The friction is ***F**_fric_* = ***ξυ*** so that ***υ*** is obtained from the force balance equation: ***υ*** = (***F**_act_* + ***F**_mem_* + ***F**_area_*)/*ξ*. The motion of the phase field *ϕ* is then determined by the advective equation *∂ϕ*/*∂t* = **−*υ*** · ∇*ϕ*. Finally, coupling the phase field equations to the reaction-diffusion equations presented in the main text, we arrive at the full equations:

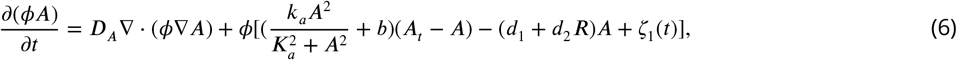

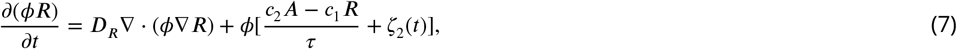

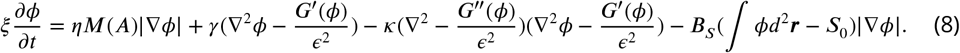

The parameters of the mechanic module, such as tension and bending, are taken from Camley et al. (2014). The parameters of biochemical module (*k_a_*, *K_a_*, etc.) are estimated from experi-mentsArai et al. (2010); Gerhardt et al. (2014), such that the minimum wave speed in simulations is approximately 0.12*μ*m/s and the wavelength is about 13*μ*m.

### Numerical details

The parameters used for numerical simulations are listed in Table 1. Equations are evolved in a region with size of *L_x_* × *L_y_*=50 × 50 *μ*m with discrete grids of n×m=256×256 and periodic boundary conditions are used. Eq.8 is discretized using the forward Euler method with *∂_t_ϕ* = (*ϕ*^(*n*+1)^ - *ϕ*^(*n*)^)/Δ*t*. Derivatives are calculated using finite difference formulas: 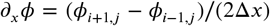 and 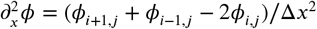, with similar equations for the derivatives in the *y*-direction. Eq.6&7 are discretized using the forward Euler scheme with 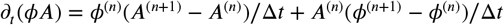. The diffusion terms ∇ · (*ϕ*·*A*) are also approximated using finite difference. The *x*-term, for example, reads 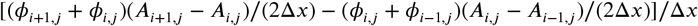. The white noise terms are simulated as Wiener processes with 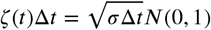. As initial condition for *ϕ*, we use a disk 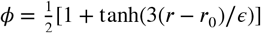, where *r*_0_ is the prescribed radius and *A* = *R* = 0. The activator and inhibitor concentration outside the boundary is 0. To implement this boundary condition, we solve Eq.6&7 only in region *ϵ*_0_ away from *ϕ* = 1/2 which is *ϕ* > *χ* = 1/2+ 1/2tanh(−3*ϵ*_0_/*ξ*) ≈ 0.0025, and leave *A* = *R* = 0 outside this region. Here, we have taken *ϵ*_0_ = 2*μ*m.

Equations are parallelized with CUDA and simulated using GPUs. Typical simulations speeds on a high-end graphics board are less than one minute for 100s of model time.

### Speed of keratocyte-like cells

The local cell boundary velocity can be approximated as 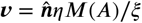. Since, for keratocyte-like cells, the front is almost flat (see main text Fig.2), the cell speed can be approximated as *υ* = *αη*/*ξ*, where *α* is the average of *M*(*A*) across the boundary: 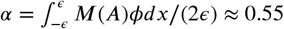. Thus, the speed of keratocyte-like cells only depends on the protrusive force and the friction but not on the tension or on the inhibitor timescale. We can verify this in simulations by changing the tension *γ* and the inhibitor’s timescale *τ* while keeping other parameters fixed. The results are shown in Fig2–Figure Supplement 1 which demonstrates that the cell speed changes little if these parameters are varied.

### Wave formation

#### Traveling waves

Our relaxation oscillator system exhibits traveling waves with a minimum speed that depends on the timescale of the inhibitor and its diffusion constant. Our model can be written as *∂_t_A* = *D*∇^2^ *A* + *f*(*A*, *R*), *∂_t_R* = *D*∇^2^ *R* + *g*(*A*, *R*), where *f*(*A*, *R*) and *g*(*A*, *R*) can be found from Eq.6&7. First we consider the case of *τ* → ∞ and *D_R_* = 0 so that the inhibitor is uniformly distributed and constant: *R* = *R*_0_. The relevant equation now is 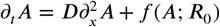. When *R*_0_ is in a proper range, there are three steady states *A* = *A*_1,2,3_, with *A*_1_ < *A*_2_ < *A*_3_ and *A* = *A*_1,3_ stable and *A* = *A*_2_ unstable. For a given *R*_0_, we seek for solutions of wave form *A*(*z*) = *A*(*x* – *ct*). The excitable version of this system has been extensively studied and for the cubic reaction term *k*(*u* – *u*_1_)(*u*_2_ – *u*)(*u* – *u*_3_) there is a stable traveling wave solution that connects *u* with *u*_1_ and *u*_3_ and has a wave speed 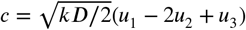 Murray (2002). Likewise, our model 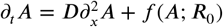, which has the same structure as the excitable system with a cubic reaction equation, has a stable traveling wave with speed *c* ≈ *w*(*A*_1_ - 2*A*_2_ + *A*_3_) which depends on *R*_0_ through *A*_1,2,3_. Here, *w* is a constant that only depends on the diffusion coefficient *D_A_* and reaction rates (cf. 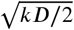 for the cubic reaction equation).

For non-zero values of *τ* and *D_R_*, the relaxation phase, characterized by the accumulation of inhibitor, sets in behind the activator’s wave front. From the above analysis, it can be deduced that a necessary condition for a stable wave front is a constant profile of the inhibitor *R* = *R*_0_ in the width of the front. For *D_R_* = 0, this can only be the case when *τ* is large so that the reaction rate of the inhibitor is slow compared to the rate of the activator. Thus, there needs to be a separation of timescales and the system has to obey relaxation dynamics. Furthermore, for *D_R_* > 0, the activator wave front is only stable if it can outrun the inhibitor’s spreading speed which is proportional to 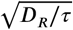. Therefore, the traveling wave is only stable if both the reaction and diffusion of the inhibitor are slow enough. In other words, stable waves will have a speed that exceeds a minimum value which depends both on *D_R_* and *τ*.

#### Transitions between migration modes

The existence of a minimum speed also indicates that if the traveling waves are bounded, the boundary has to move fast enough to create space for the wave front. In our model, we can tune *η* to change the boundary mobility (cf. Eq. 4 main text). For small values of *η*, the speed of the boundary will be smaller than the minimum speed required for stable wave propagation and traveling waves that hit the boundary will stall. For larger values of *η*, exceeding the first transition in the phase diagram of Fig. 2, the speed of the boundary is large enough to accommodate the traveling activator wave.

One important consequence of the above reasoning is that there will be a transition from non-motile to motile cells as the protrusive force *η* is increased. For small value of *η* the center of mass of the cell will not show significant movement, while for larger values of *η* the transient membrane protrusions will result in net center-of-mass motion. In simulations, we change *η* and monitor the cell’s center-of-mass speed *υ_CM_*, as shown in Fig2–Figure Supplement 2 a. As initial condition, we can either perturb the homogeneous solution in *A* and *R* with noise or we can choose an asymmetric distribution of *A* and *R*. For both initial conditions, the speed shows a subcritical bifurcation at a critical value of *η*_*c*,1_, above which a non-zero cell speed emerges. The non-zero cell speed at the bifurcation point corresponds to the minimum wave speed *c*_min_ since at this point *c*_min_ = *αη*_*c*,1_/*ξ*. In simulations, we can change *c*_min_ by changing the inhibitor’s diffusion coefficient *D_R_* and the timescale *τ*. As expected, larger *D_R_* and smaller *τ* leads to larger *c*_min_, and consequently larger *η*_*c*,1_ (Fig2–Figure Supplement 2b&c).

#### The role of tension in morphology

Tension is important to maintain the unidirectional movement of keratocyte-like cells. In our model, at the two lateral ends of the traveling wave the morphology is determined by a balance between the protrusive force and the tension which is determined by the local curvature and the parameter *γ*. When the protrusive force increases, a larger membrane curvature is necessary to balance the protrusive force, resulting in a flatter front and a decreased front-back distance (see Fig2–Figure Supplement 3 a). If *γ* is not large enough, the traveling wavefront can have a turning instability: the cell will no longer migrate along a straight path and will make a turn Camley et al. (2017). An example of this instability and the resulting motion is shown in Fig2–Figure Supplement 3 b where we decrease the surface tension by a factor of 2 (from *γ*=2pN to *γ*=1pN). Similar to Camley et al. (2017), the cell can also be destabilized by increasing the diffusion of the activator and inhibitor. An example of a simulation showing this can be found in Fig2–Figure Supplement 3 c.

#### Parameter variations

We have examined how the model results change when certain parameters are varied. For example, we can extend the (*η*, *r*) phase space to larger values of *r* as shown in Fig2–Figure Supplement 4 a. For cell sizes that are beyond ones observed in experiments, we And that the critical protrusive force *η*_*c*,2_ saturates. For these large values of *r*, a dominant wave forms at the front of the cell and new waves that are generated at the back of the cell are not strong enough to break this dominant wave’s persistency. The cell will move persistently in the direction of the dominant wave, with smaller waves repeatedly appearing at the back, as shown in the snapshot presented in Fig2–Figure Supplement 4 a. For keratocyte-like cells with a single wave, *d* will saturate to the wavelength *λ* ≈ 13*μ*m, as shown in Fig2–Figure Supplement 4 b.

The transition from amoeboid-like cell to keratocyte-like cell is determined by the front-back distance *d* and occurs when *d* is smaller than the spatial scale of the wave *λ*. To verify this, we vary parameters that either affect *d* or *λ*. First, we reduce the timescale of the inhibitor *τ* to half the value reported in Table 1 (*τ*_0_) which leads to a decrease of *λ* and should lead to larger values of *η*_*c*,2_. Our simulation results, presented in Fig2–Figure Supplement 4 c, show that the critical value of the protrusive strength is indeed larger than the one for the default parameters (blue line vs. black line). Next, we increase the value of the membrane tension to *γ* = 2*γ*_0_. Our simulations reveal that for increasing values of the membrane tension the transition occurs for larger values of *η* (Fig2–Figure Supplement 4 c, green line). This can be understood by realizing that an increase in the membrane tension will reduce the cell’s deformability. Therefore, the curvature of the cell’s front will decrease, which will increase the front-back distance and thus the critical protrusion strength. Finally, we double the value of the friction coefficient *ξ* which should, according to Eq.4 in the main text, lead to a decrease in the membrane speed. Since the biochemical wave speed *c* is unchanged, the transition from amoeboid-like to keratocyte-like motion should occur for larger values of *η*. This is confirmed in our simulations, which show that the bifurcation line for *ξ* = 2*ξ*_0_ occurs for larger values of *η* (yellow line in Fig2–Figure Supplement 4 c).

#### Varying area conservation

Real cells are three-dimensional objects in which the changing of the basal surface area will be compensated by the morphological changes away from the substrate. Our model represents the cell as a two-dimensional object and therefore includes an area conservation term. Making the strength of this area conversation large in simulations allows us to define and sample the (*η*, *r*) phase space (Fig. 2b). To determine how this area conservation term affects the observed dynamics, we reduce the area conservation parameter from *S_B_* = 10 to *S_B_* = 0.1. As shown in Fig2–Figure Supplement 5, all migration modes are still present. For small values of *η*, cells are oscillatory and, compared to large values of *S_B_*, exhibit measurable oscillations in cell size (Fig2–Figure Supplement 5 a-c). Furthermore, increasing the value of the protrusive strengths results in amoeboid-like motion while even larger values of *η* lead to keratocyte-like cells (Fig2–Figure Supplement 5 d).

### Excitable model

Our biochemical model is based on relaxation oscillation dynamics. However, it is straightforward to consider an excitable version of the model. For this, we take *c*_2_ = 30, *σ* = 0.1 and keep all the other parameters same as listed in Table S1. For the excitable version of our biochemical module, we find similar migration mode transitions as found in the main text as well as a qualitatively similar phase diagram (Fig2–Figure Supplement 5 a and b). Specifically, we also find a nonmotile, amoeboid-like, and a keratocyte-like mode (Fig2–Figure Supplement 5 a). In the excitable case, however, perturbations are required to initiate waves and movement. As a consequence, the patterns of activator are more noisy than for the oscillation model and the non-motile cells do not exhibit oscillatory dynamics. Finally, the transition between nonmotile cells and amoeboid-like cells is also subcritical (Fig2–Figure Supplement 5 c).

### Experiments

Wild-type AX2 cells were transformed with the plasmid expressing limE-delta-coil-GFP. Cells were kept in exponential growth phase in a shaker at 22°C in HL5 media with hygromycin (50*μ*g/mL). On the day before the experiment, cells were diluted to low concentration (1-2×10^5^ cells/mL) to stop the exponential growth. After 15h-18h, cell concentration reached 2-5×10^5^ cells/mL and 10^5^ cells were plated in a 50mm round chamber with glass bottom (WillCo). After 15min, cells attached to the substrate and HL5 was replaced with 7mL DB (5 mM Na_2_HPO_4_, 5 mM KH_2_PO_4_,200*μ*M CaCl_2_, 2 mM MgCl_2_, pH6.5).

Specified amount was diluted in 500*μ*L of DB and added to the sample using a pipette at 6h of starvation.

Differential interference contrast (DIC) images were taken every 30s in six fields of view across the sample using a 10x objective from 5h45 to 6h30 after beginning of starvation. Cell centroids, area, minor and major axis were tracked using Slidebook 6 (Intelligent Imaging Innovations). Statistical analysis of trajectories was performed in MATLAB (2018a; The Mathworks). Angles between two successive positions, separated by 30s, were computed. The standard deviation of this angle was computed for 5 consecutive pairs and keratocyte-like cells were defined by having a standard deviation less than 25 degrees. Experimental data presented before latrunculin B are for cells between 5h45 to 6h of starvation, whereas effect of the drug is quantified on cells from 6h15 to 6h30. Speeds were measured using a time interval of 1min. For each concentration, data were collected on three different days. P values were computed with the Wilcoxon rank sum test. Fluorescent images (488-nm excitation) were captured with a 63x oil objective using a spinning-disk confocal Zeiss Axio Observer inverted microscope equipped with a Roper Quantum 512SC cameras.

### Movies

Movie S1: simulation results for *r* = 8*μ*m and *η* = 2 pN. Movie S2: simulation results for *r* = 8*μ*m and *η* = 4 pN. Movie S3: simulation results for *r* = 8*μ*m and *η* = 11 pN. Movie S4: experimental results with 2*μ*M Latrunculin B added at time 1h45min.

---

**Figure 2–Figure supplement 1.**
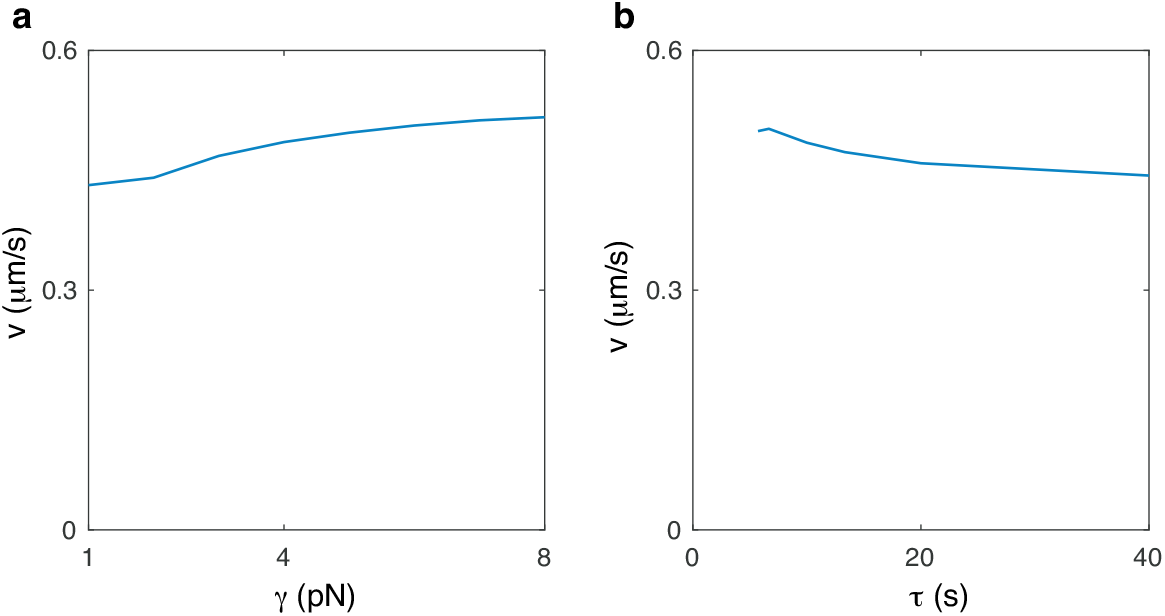
Speed of keratocyte-like cells as a function of the surface tension (a) and the timescale of the inhibitor (b). Parameters are as in Table S1 with *r* = 8*μ*m.

**Figure 2–Figure supplement 2.**
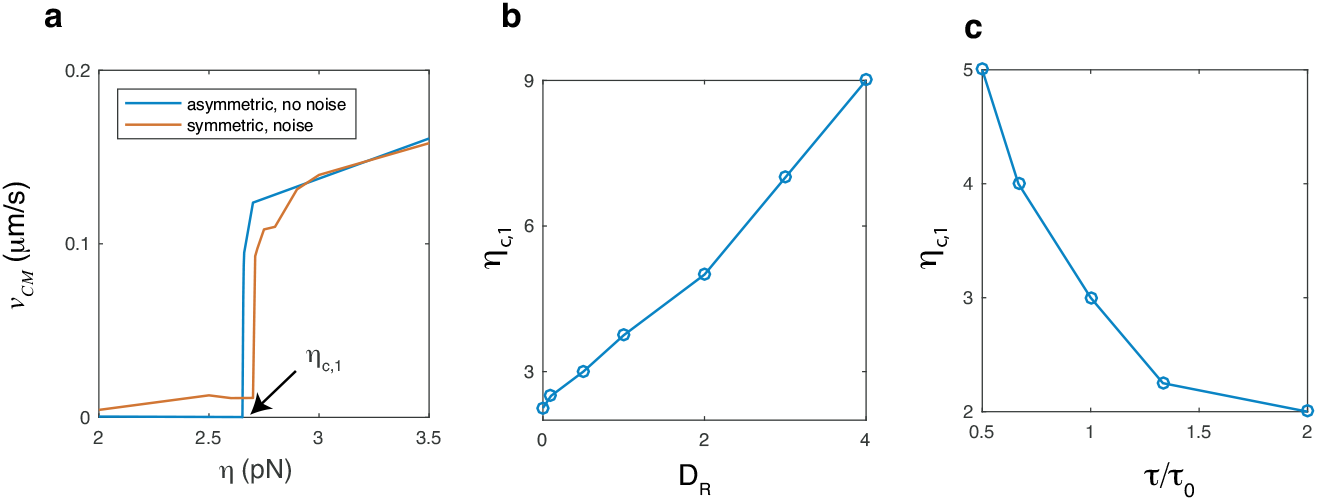
(a) Speed of the center of mass of a cell as a function of protrusion strength *η*. The red curve represents results from initial conditions where noise is added to a homogeneous *A* and *R* field while the blue curve correspond to simulation in which the initial activator is asymmetric. Cells become non-motile at a critical value of protrusion strength, *η*_*c*,1_. The critical value of protrusion strength as a function of, (b), inhibitor’s diffusion constant *D_R_*, and (c), the timescale of the inhibitor *τ*, rescaled by the default value *τ*_0_ = 10s. Remaining parameters are as in Table S1 with *r* = 8*μ*m.

**Figure 2–Figure supplement 3.**
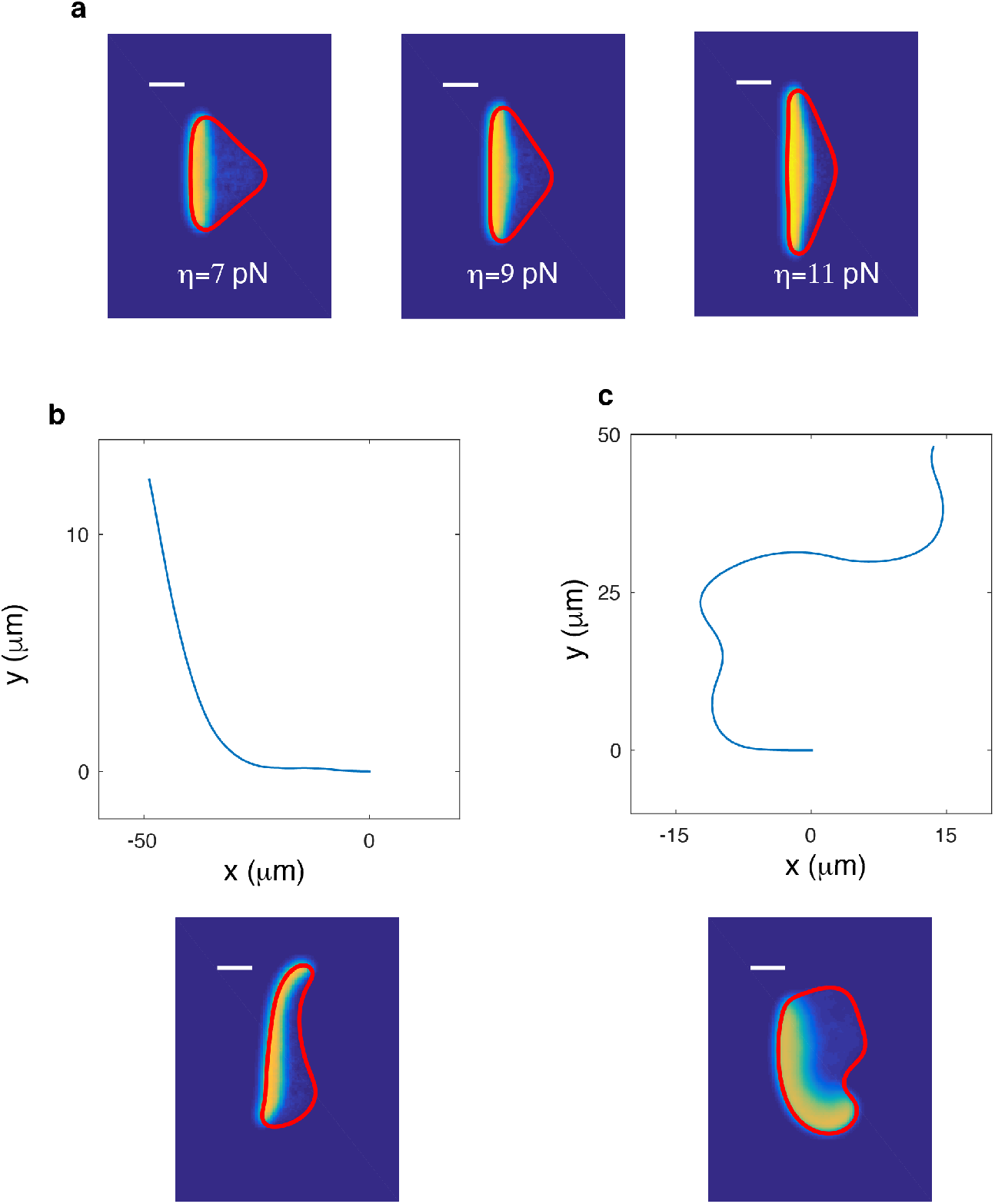
(a) Increasing the protrusive force *η* will result in flatter front and a decreased front-back distance. (b) Snapshot (bottom) and corresponding trajectory (top) of a cell that undergoes a turning instability upon the reduction of surface tension from *γ*=2pN to *γ*=1pN. Parameters are taken from Table S1 with *r* = 6*μ*m. (c) As in (b), but now after an increase in *D_A_* and *D_R_* to *D_A_* = *D_R_*=2 *μ*m^2^/s and *r* = 8*μ*m.

**Figure 2–Figure supplement 4.**
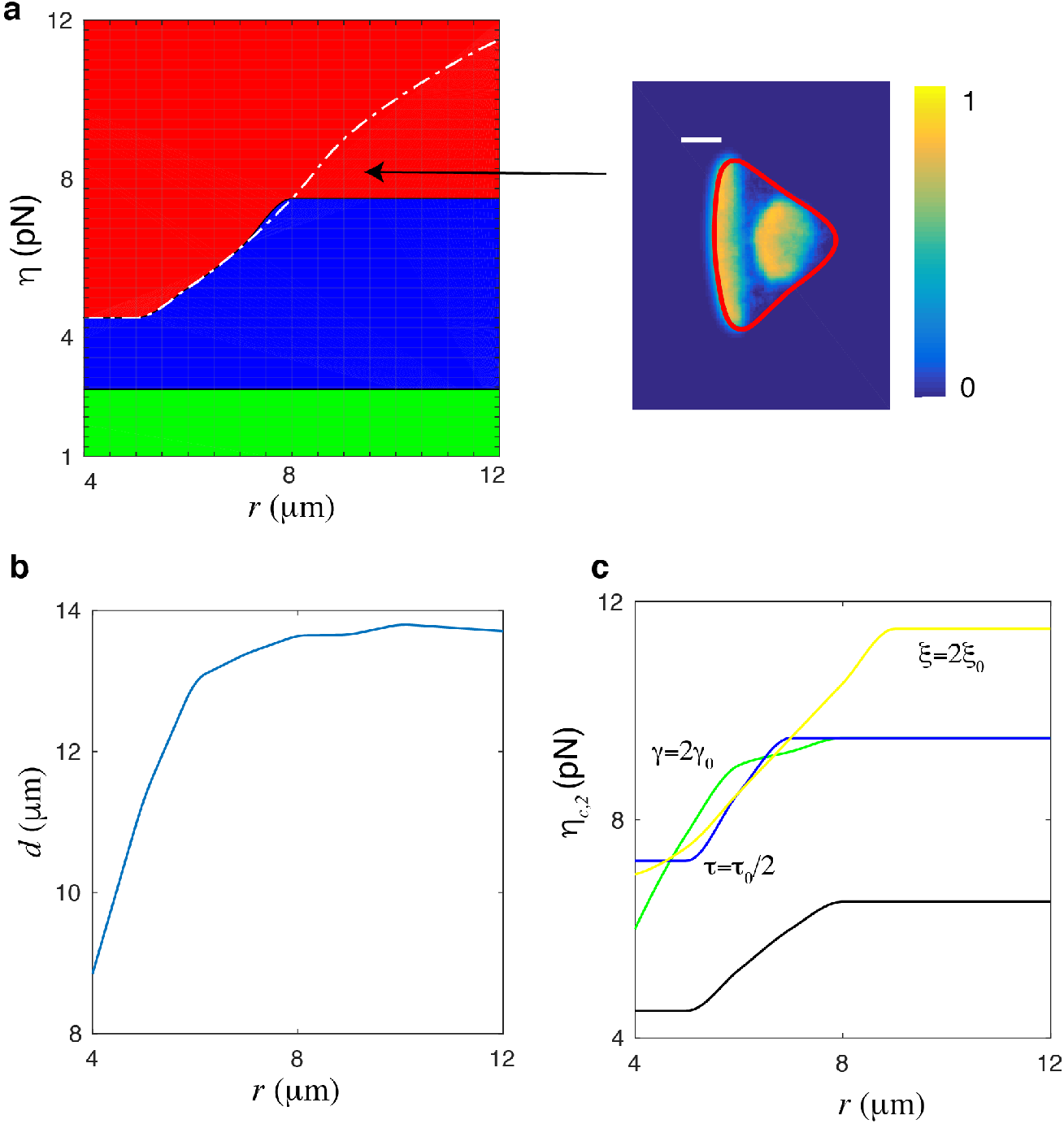
Parameter variations in the model. (a) Phase space extended to maximum cell area ~ 450*μ*m^2^, corresponding to *r* = 12*μ*m. Green: oscillatory cell; blue: amoeboid-like cell; red: keratocyte-like cell. The dashed white line corresponding to the separation line *d* = *λ* above which the keratocyte-like cells exhibit a single wave and below which keratocyte-like cells move unidirectional but with small waves repeatedly appearing at the back of the cell. (b) The keratocyte-like cell’s front-back distance *d* along the white dashed line. The saturation value is the wavelength *λ* ≈ 13*μ*m. (c) The transition line of amoeboid-like cell to keratocyte-like cell under different parameters.

**Figure 2–Figure supplement 5.**
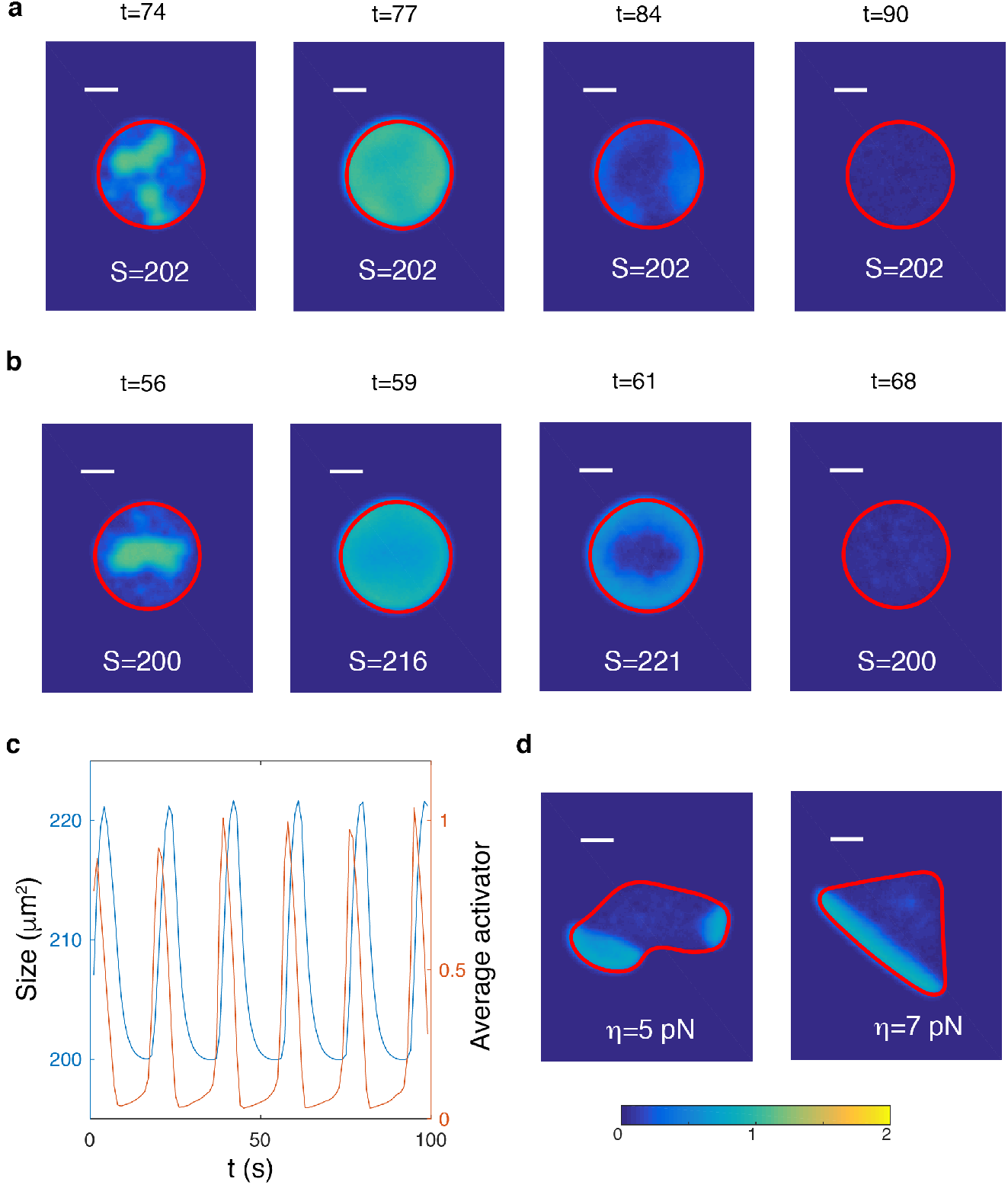
Oscillatory cells for strong (a, *B_S_* = 10) and weak (b, *B_S_* = 0.1) area conservation. Parameters are as in Table 1 with *η* = 2.5 and *r* = 8*μ*m. (c) Cell size (blue) and average activator concentration (red) as a function of time for the oscillatory cell in panel b. (d) Left, example of amoeboid-like cell for weak area conservation (*B_S_* = 0.1); right, example of keratocyte-like cell for weak area conservation (*B_S_* = 0.1). The colors represent the activator concentration, as indicated by the color bar. Scalebar=5 *μ*m.

**Figure 2–Figure supplement 6.**
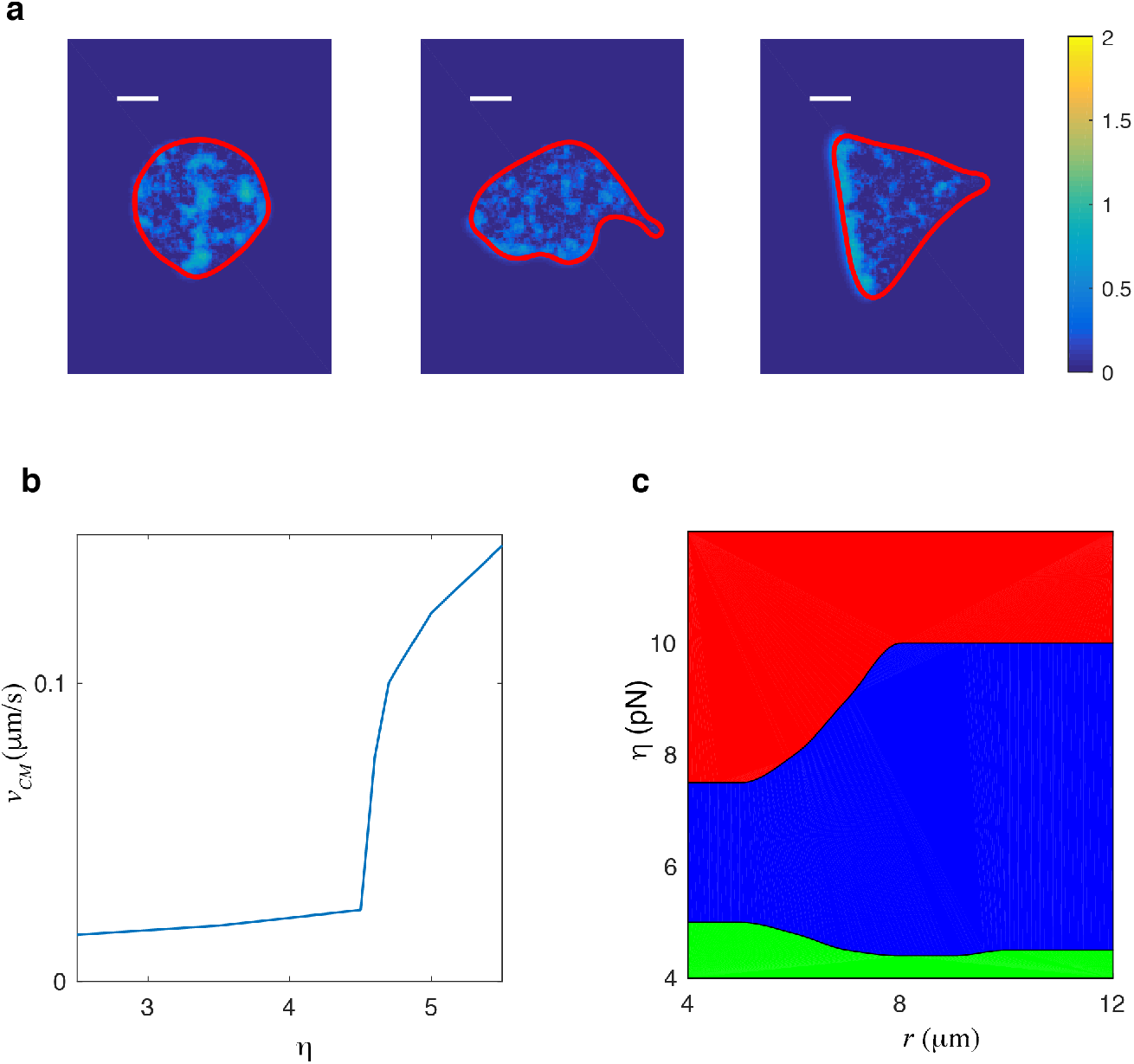
Excitable dynamics can reproduce identical qualitative results. (a) Snapshots of the three different migration modes for an excitable version of the model. Left panel shows an nonmotile cell (*η* = 2), middle panel shows an amoeboid-like cell (*η* = 4), and the right panel displays a keratocyte-like cell (*η* = 10). (b) Speed of the center of mass as a function of critical protrusion strength. As Fig. S2a, the bifurcation is subcritical. (c) Phase diagram (*η-r* space) with green corresponding to nonmotile cells, blue representing amoeboid-like cells, and red corresponding to keratocyte-like cells. Scalebar=5 *μ*m.

**Figure 3–Figure supplement 1.**
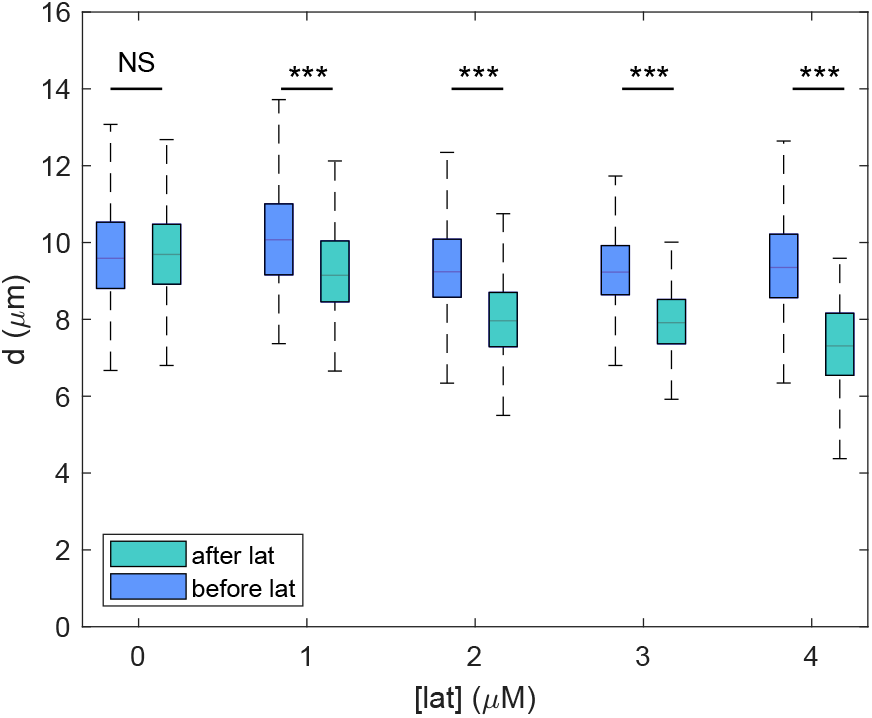
The front-back distance of keratocyte-like cells before and after latrunculin exposure as a function of concentration.

**Figure 3–Figure supplement 2.**
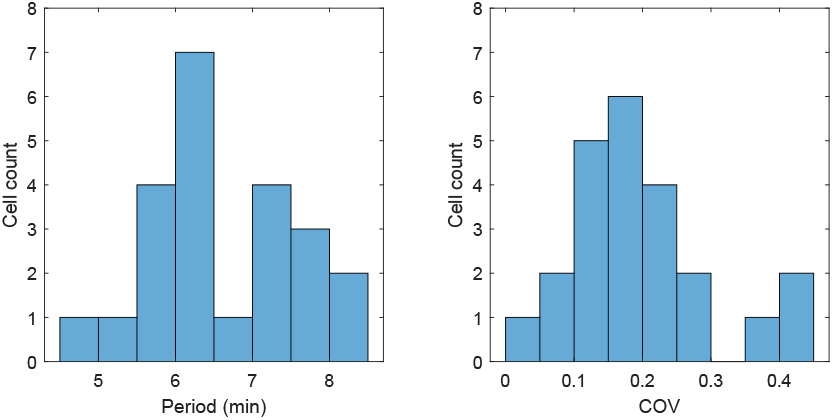
The average period and coefficient of variation COV (ratio of the standard deviation and the mean) for area oscillations in oscillating cells after the exposure to 4*μ*M latrunculin (N=23).

